# The genomic sequence and comparative genomic analysis of cultivated passion fruit(*Passiflora edulis* L.)

**DOI:** 10.1101/522128

**Authors:** Yanyan Wu, Qinglan Tian, Jieyun Liu, Yongcai Huang, Weihua Huang, Xiuzhong Xia, Haifei Mou, Xinghai Yang

## Abstract

Cultivated passion fruit is a fruit tree widely cultivated in southern China, but little is known about its genomics, which seriously restricts the molecular genetics research of passion fruit. In this study, we analyzed the 165.7Mb representative genome sequences. The results showed that the passion fruit genome contained a large number of simple sequence repeats (SSR). Compared to the cassava and peach genomes, the passion fruit genome has 23,053 predicted genes. These genes can be aligned to 282 plant genomes. GO annotation indicated that these genes are involved in metabolic pathways of carbohydrates, organic acids, lipids and other molecules. KEGG pathway enrichment assigned these genes into five major categories and 19 secondary functions. Cluster analysis of gene families showed that 12,767 genes could be clustered into 9,868 gene families and 291 unique gene families. On the evolutionary relationship, the passion fruit is closely related to *Populus trichocarpa* and *Ricinus communis*, but the rate of evolution is slower. In summary, this genomic analysis result is informative, and will facilitate the future studies on gene functions of passion fruit.

## 1. Introduction

There are more than 530 species of passion fruit, and the most widely cultivated species is *Passiflora edulis*, which belongs to the *Theoideae* suborder, *Passifloraceae* family, and *Passiflora L.* genus [1]. Passion fruit has really high contents of nutrition, including sugar, fat, protein, vitamins and mineral elements [2,3]。

In eukaryotes, the genome is the entire genetic material of a single set of chromosomes in the species. Each cell of a plant contains three distinct genomes: nuclear genome, mitochondrial genome, and plastid genome. Currently, studies are mainly focused on the nuclear genome. Chromosomes are gene carriers, and the gene functions are closely related to the structural components on chromosomes. Genome sequencing can help us better understand the functions and evolution of plant genes.Currently, the genome sequencing study on passion fruit is still focused on the development of molecular markers. Cerqueira-silva et al. [4] developed 69 pairs of SSR primers using two passion fruit genome microsatellite-enriched libraries. Santos et al. [5] used BAC end sequencing method to obtain 6,194,248 bp of passion fruit genome data, in which 669 microsatellite sequences were found, with an average of one SSR per 9.25 kb genome sequence. Later, Araya et al. [6] developed 816 pairs of SSR primers in the structural and functional regions using parts of the passion fruit genome sequence. The results showed that 53.2% of SSR primers were polymorphic. Recently, Costa et al. [7] sequenced the cDNA of Xanthomonas infected passion fruit, and developed the functional SSR and SNP markers.

With the rapid development of High-throughput sequencing, nearly 200 plants have been sequenced. In May 2017, the Beltsville Agricultural Research Center performed genome-wide sequencing on passion fruit CGPA1 using Illumina GAII sequencing technology, and assembled the sequencing results to the Scaffold level. However, they did not conduct genome analysis on these results. In this study, we performed genome annotation and comparative genomic analysis on passion fruit genome. Our results will facilitate the further studies on molecular mechanisms of passion fruit, and also provide references for the scientific development and efficient utilization of passion fruit.

## 2. Materials and Methods

### 2.1. Genomic Sequence of Passion Fruit

The passion fruit genome was uploaded to NCBI (https://www.ncbi.nlm.nih.gov/assembly/GCA_002156105.1/#/st) by the Beltsville Agricultural Research Center.

### 2.2. Genome Annotation of Passion Fruit

Identification of autonomous DNA transposon: The known autonomous DNA transposons in plants, such as Arabidopsis, were collected from public databases (Swiss-Prot and Repbase). Then, the transposons in passion fruit were identified by the software detectMITE [8].

Gene structure prediction: Homologous prediction was conducted by comparing the protein coding sequence of a known homologous species with the genomic sequence of a new species (the number of homologous species is no more than 5). The gene structures of new species were predicted by softwares such as BLAST (http://blast.ncbi.nlm.nih.gov/Blast.cgi), GeneWise [9], etc. De novo prediction used the software depending on statistical characteristics of genomic sequence data to predict gene structure. The commonly used software includes Augustus [10], Glimmer HMM [11], SNAP (http://homepage.Mac.com/iankorf/), etc. After performing the gene structure prediction, the results were combined with the transcriptome alignment data; then, these data were integrated by the EVidenceModeler software (http://evidencemodeler.sourceforge.net/) to generate a non-redundant, more complete gene set. Finally, the EVM annotation results were corrected using PASA (http://pasa.sourceforge.net/) and the transcriptome assembly data. The information such as UTR and variable cutting sites was added to obtain the final gene set.

Gene function annotation: The gene set obtained by gene structure annotation was compared with a known protein database by comparison software, in order to obtain the gene function information. The commonly used protein databases include SwissProt (http://www.uniprot.org/), KEGG (http://www.genome.jp/kegg/), InterPro (https://www.ebi.ac.uk/interpro), NR (ftp://ftp.ncbi.nlm.nih.gov/blast/db/) and GO (http://www.geneontology.org/).

### 2.3 Gene Family and Phylogenetic Tree

#### Gene family identification

The software OthoMCL[12] was used. The default e value was 1e−5 and the expansion coefficient was 1.5.

#### Phylogenetic analysis

The software MUSCLE [13] was used to compare different gene families. The sequence alignment results went through jModelTest/ProTest [14] software to find the optimal sequence substitution model. Then, the phylogenetic tree of 9 species was constructed by PhyML software [15] using the maximum likelihood method.

## 3. Results

### 3.1. Assembly of Passion Fruit Genome

The research group at Beltsville Agricultural Research Center used Illumina GAII technology to sequence the passion fruit CGPA1 genome. The average sequencing depth was 4.5×, with 225,293,527 reads in total. Finally, 165,656,733 bp of passion fruit genome sequence was obtained, with 235,883 Contig (Contig N50 was 1,303 bp, Contig L50 was 30,212 bp) and 234,012 scaffolds (Scaffold N50 was 1,311 bp, Scaffold L50 was 30,081 bp). The GC content of the genome was 38.6%.

### 3.2. Repeated Sequence Annotation

The SSR Search software [16] and homologous annotation were used to annotate the repeated sequences in passion fruit genome. The results showed that there were 428,294 full-type SSR and 1,544,549 incomplete- and composite-type SSR [6]. For transposons, there were 59 *Mutator* transposons, 41 *EnSpm* transposons, 49 *hAT* transposons, 221 *PIF* transposons, and 2 *MLE* transposons.

### 3.2. Gene Annotation and Functional Enrichment Analysis of Passion Fruit Genome

Genetic structure prediction was conducted using homologous prediction and De novo prediction. Using BLAST, GeneWise, and other alignment softwares, the genomic sequence of passion fruit was compared with the coding sequences of known homologous species *Manihot esculenta* [17] and *Prunus persica* [18] to predict the gene structures in passion fruit genome. These prediction results were then combined with the transcriptome alignment data, and all the gene sets predicted by different methods were integrated by the EvidenceModeler software to generate a non-redundant and more complete gene set. Finally, the EVM annotation results were corrected using PASA and transcriptome assembly results. The information such as UTR and variable cutting sites were added, and 23053 genes were eventually predicted.

The gene set obtained by gene structure prediction was blasted in NR, SwissPort, KEGG, InterPro, Pfam and GO databases, and the gene annotation information was shown in Table 1. In KEGG database, the passion fruit genome had 16,835 genes annotated. The gene length was 61-6994 bp, with an average of 670 bp. The total length of annotated genes was 11,784,169 bp, accounting for 7.1% of the whole genome. The predicted passion fruit genes can be mapped to the genomes of 282 plant species. Among these genes, 3,015 of them were aligned to the *Populus trichocarpa* genome, 2058 genes were mapped to the *Jatropha curcas* genome, 1,644 genes were aligned to the *Ricinus communis* genome, 630 genes were mapped to the *Theobroma cacao* genome [19], and 572 genes were aligned to the *Vitis vinifera* genome [20].

**Table 1.**
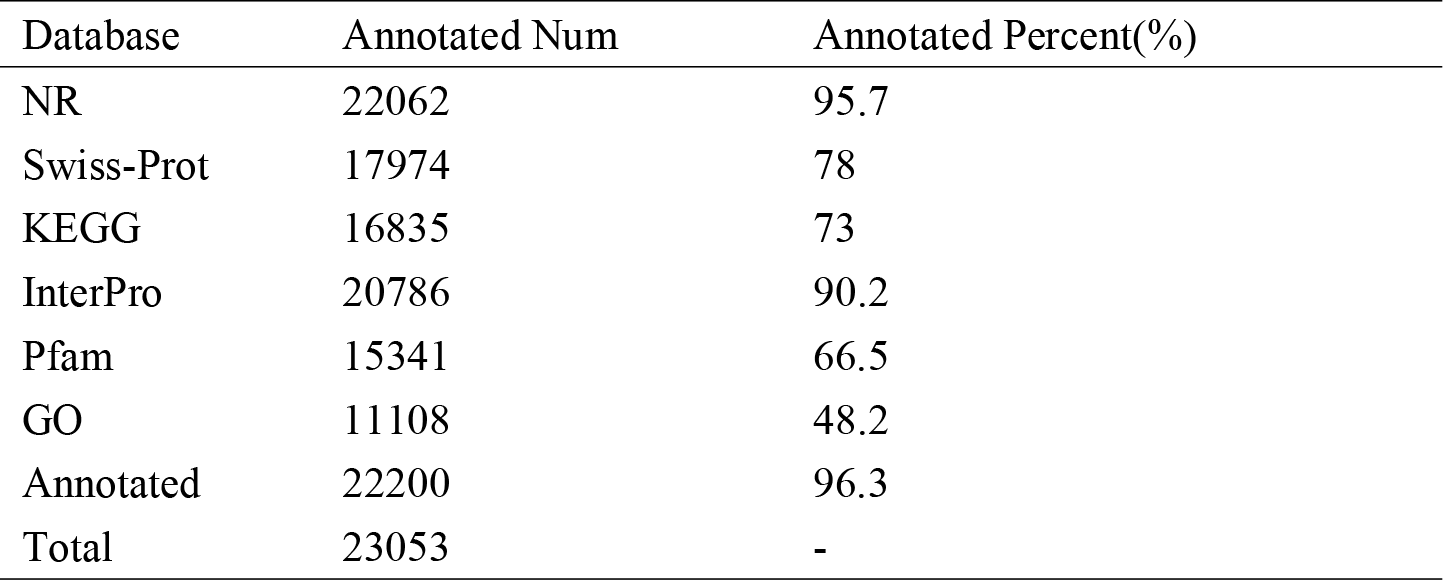
The annotation cultivated passion fruit genome.

GO analysis was used to classify the functions of annotated genes into categories of Biological process, Cellular component and Molecular Function; then, these functions were further refined into 41 secondary functions (Figure 1). In the Biological process category, there were more genes involved in cellular process (GO: 0009987) and metabolic process (GO: 0008152), accounting for 4,689 and 5,047 genes, respectively; in the Cellular component category, more genes were involved in cell part (GO:0044464) and cell (GO: 0005623), both of which included 1,542 genes; in the Molecular Function category, the catalytic activity (GO: 0003824) and structural molecule activity (GO: 0005198) included more genes, accounting for 5,018 and 5,595, respectively. Since passion fruit has a pleasant aromatic odor and has high contents of sugar, fat, protein, vitamins and minerals [2,3], we focused our study on the metabolic processes of carbohydrates, organic acids, lipids, etc., and found that 1,356 genes were involved in the metabolism of aromatic compounds.

**Figure 1.**
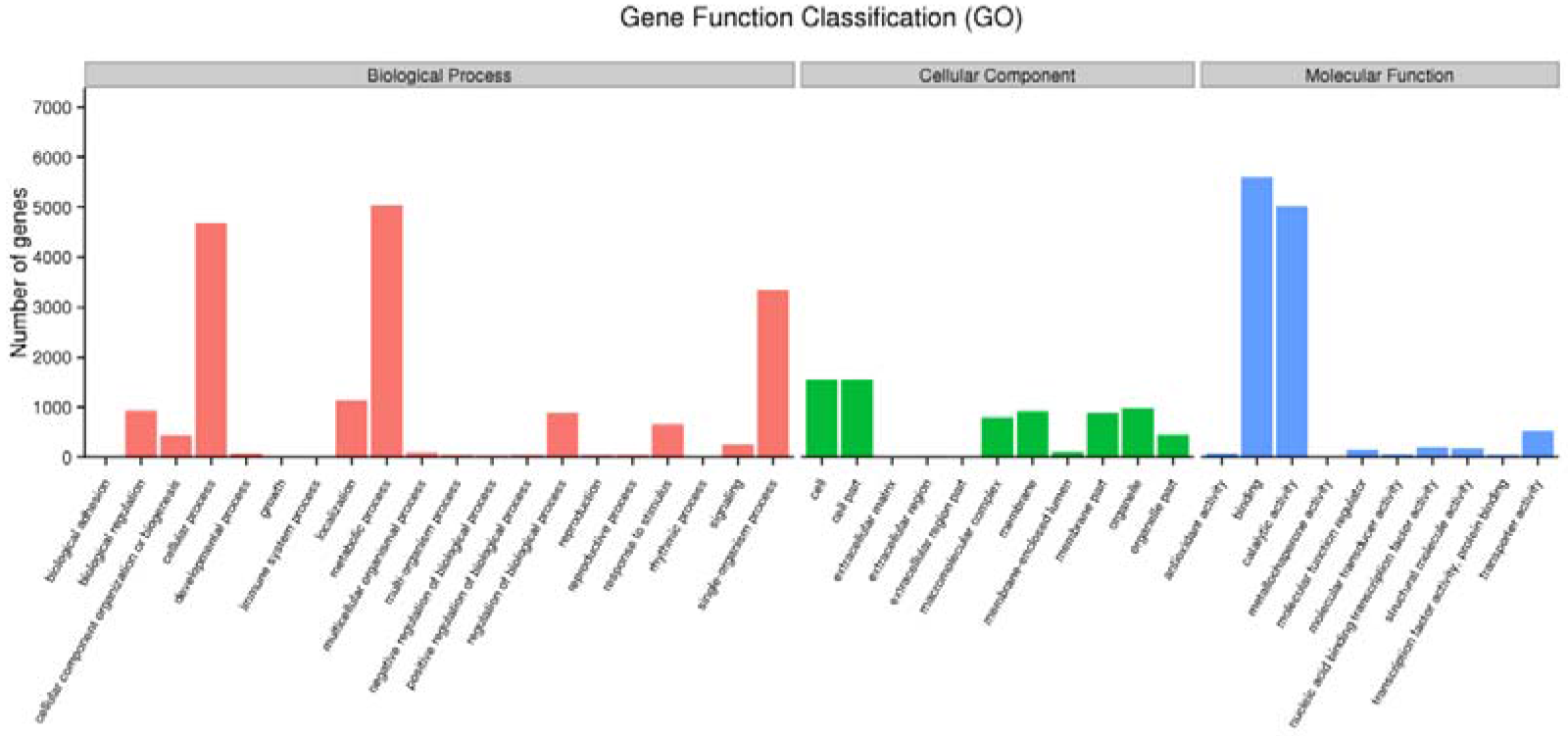
GO function analysis of the annotated genes.

In living organisms, different genes were coordinated to perform biological functions. The same actions between different genes form a pathway, and the pathway-based analysis is helpful for further interpreting the gene functions. KEGG database was used to analyze the gene pathways, and the results showed that the gene pathways were divided into five categories according to the pathway type (Figure 2): A: Cellular Processes; B: Environmental Information Processing; C: Genetic Information Processing; D: Metabolism; E: Organismal Systems. These five categories can be subdivided into 19 secondary functional classes. Among the 1,1325 genes, 61.6% were associated with metabolic pathways, and the largest group was related to carbohydrate metabolism. Glucose, sucrose, starch and cellulose are the main forms of carbohydrates. Studies have shown that passion fruit is rich in sugars and fats [2,3]. In the passion fruit genome, there were only 570 genes involved in environmental adaptation, suggesting that passion fruit may be less capable to resist biological or non-biological stresses.

**Figure 2.**
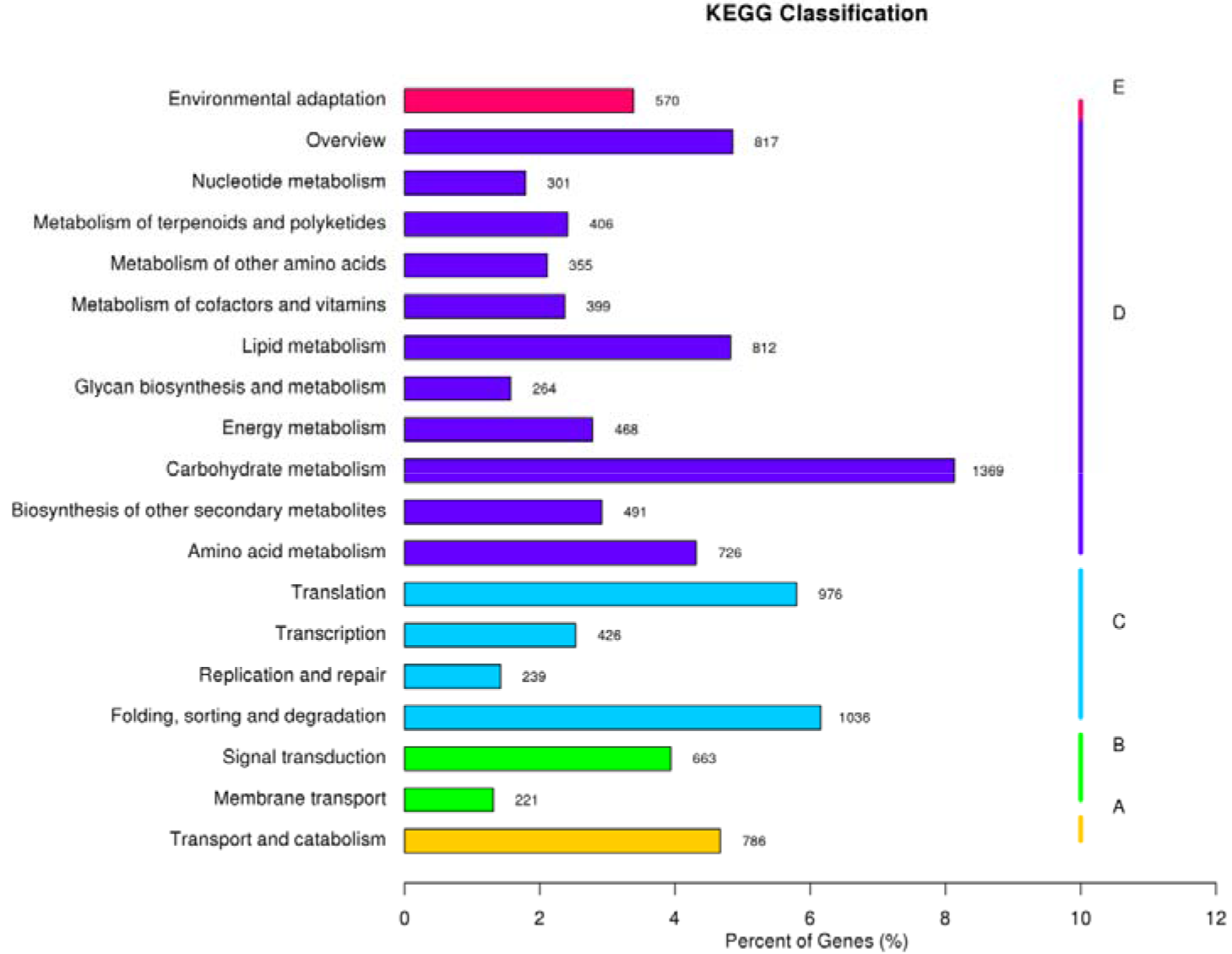
Pathway classification of the annotated genes.

### 3.3. Gene Family and Phylogenetic Analysis

Based on the passion fruit genome annotation results and the previous studies [1,5], we performed gene family analysis using another nine species, which were *Actinidia chinensis* [21], *Theobroma cacao* [19], *Vitis vinifera* [20], *Arabidopsis thaliana* [22], *Populus euphratica* [23], *Prunus persica* [18], *Ricinus communis* [24], and *Oryza sativa L. ssp. japonica* [25]. The number of aligned genes in each species is shown in Table 2. Via cluster analysis of gene families, we found 12,767 genes of passion fruit could be clustered into 9868 gene families, with an average of 1.29 genes per family. Moreover, there were 291 gene families that were unique for passion fruit (Figure 3).

**Figure 3.**
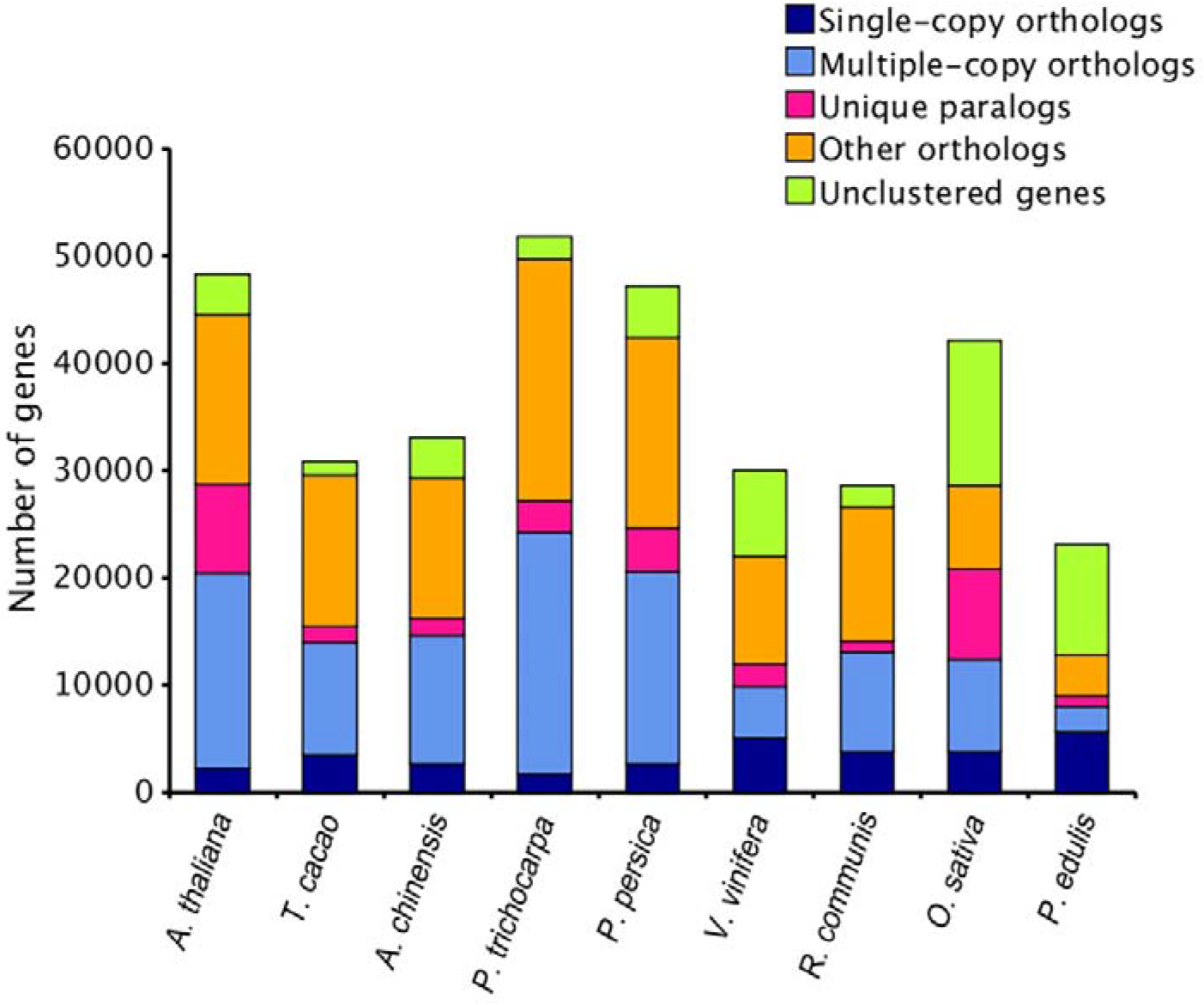
Gene family clustering in 9 species.

**Table 2.**
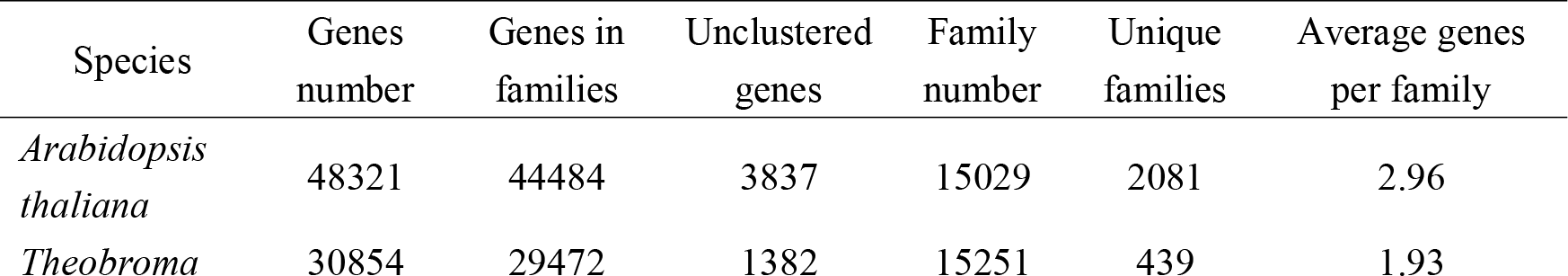
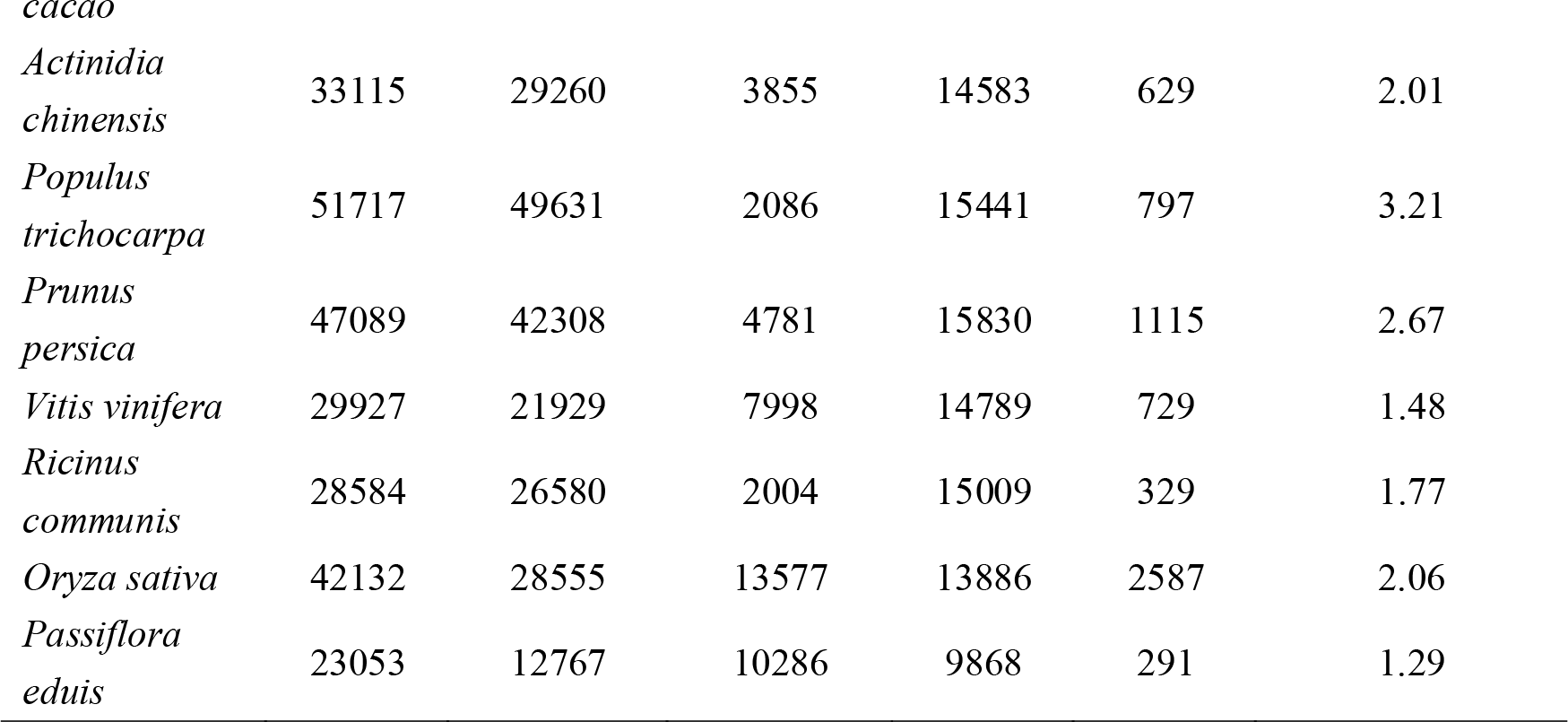
Genes used for gene family clustering in nine species.

Referring to the study from Santos et al. [5], we selected the genomes from *Actinidia chinensis* [21], *Theobroma cacao* [19], and *Vitis vinifera* [20] to perform homologous analysis with the predicted genes of passion fruit (Figure 4). The results showed that *Theobroma cacao* had the most homologous genes with passion fruit. Using *Oryza sativa L. ssp. japonica* genome as the reference, we also did phylogenetic analysis on the nine species with homologous genes (Figure 5). The cluster analysis showed that the monocots were clearly separated from the dicots. Also, passion fruit was evolutionarily closer to *Populus trichocarpa* and *Ricinus communis*, but the evolution rate was slow.

**Figure 4.**
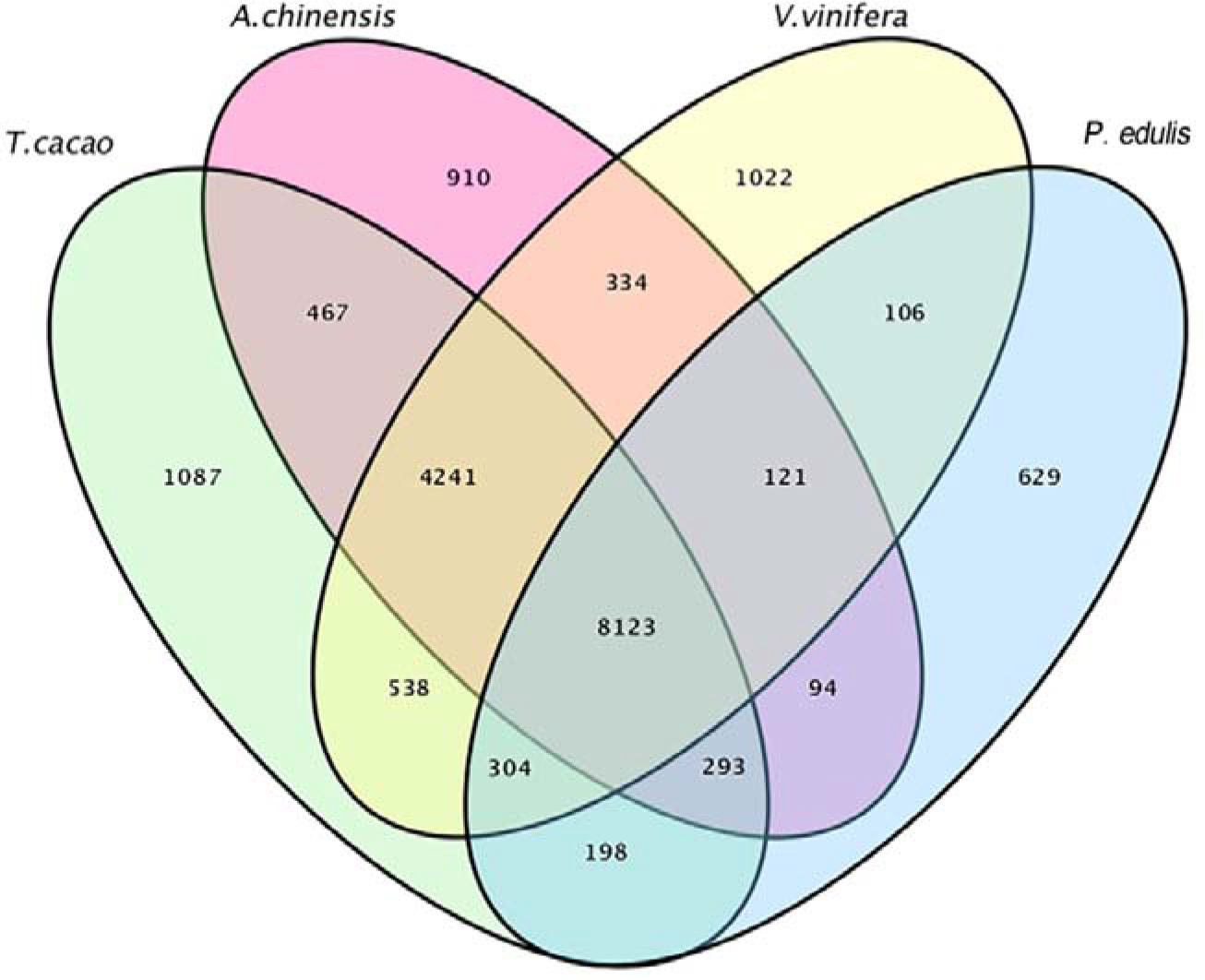
Homology analysis in 4 species.

**Figure 5.**
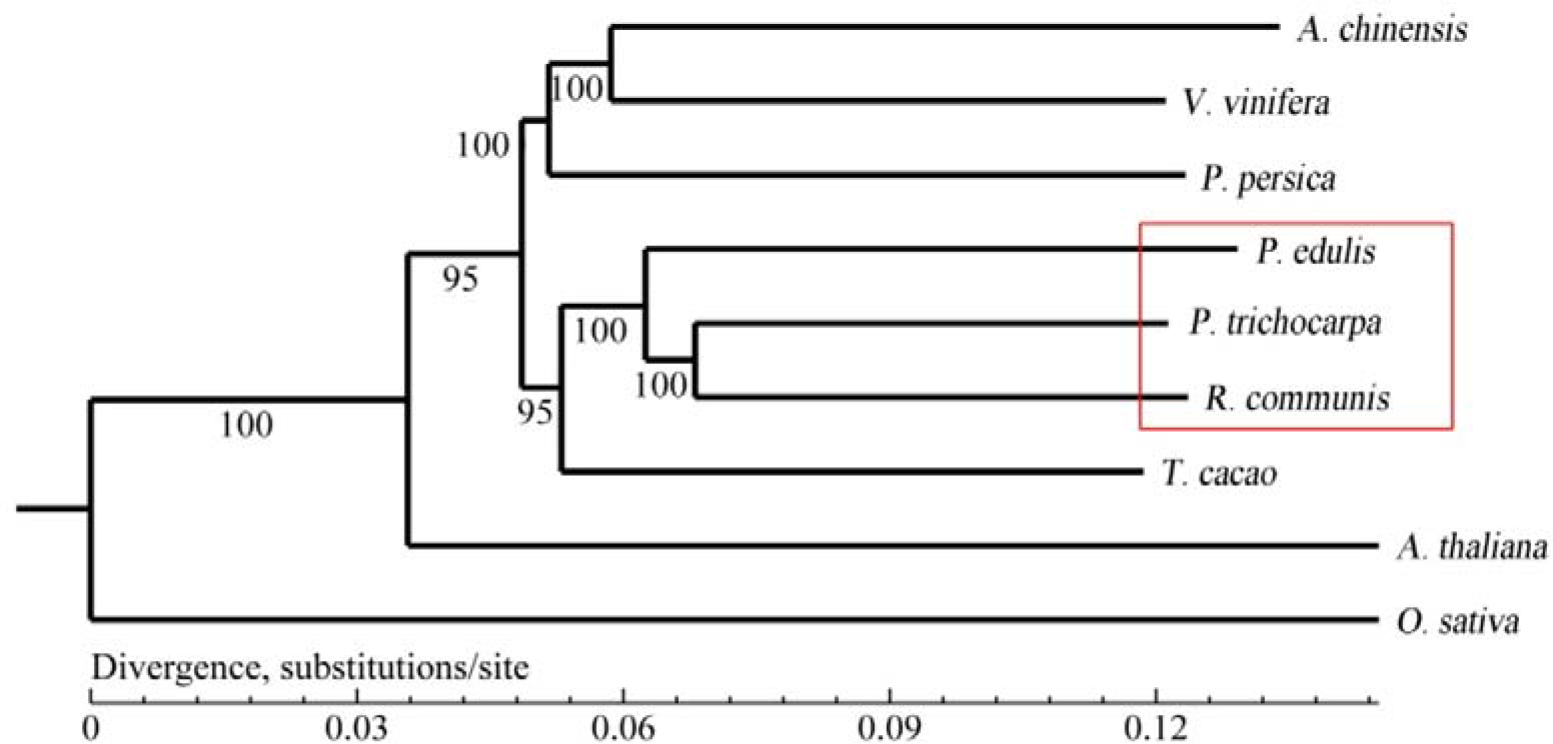
Phylogenetic analysis of 9 species.

## Discussion

The passion fruit genome is rich in repetitive elements, which can be used to develop molecular markers. In our previous study, We identified 13,104 perfect SSRs in the 165.6 Mb of cultivated passion fruit genome. Then we developed 12,934 pairs of SSR primers using a full-type SSR, and the SSR marker showed good polymorphism [16]. According to the different transposon vectors, transposons can be divided into two types: retrotransposons (Class I) and DNA transposons (Class II). The former is mediated by RNA and the latter is mediated by DNA. MITEs (Miniature Inverted Repeat Transposable Elements) are a special class of non-autonomous DNA transposons that are distributed in high-copy form in the genome of plants. The MITEs transposon marker developed by MITEs can only amplify two bands in general, and the PCR product can be efficiently isolate by conventional agarose gel electrophoresis, so the marker is highly efficient and co-dominant molecular marker. We used softwares to identify the MITEs transposon of the cultivated passion fruit genome, and obtained 372 transposons and their flanking sequences, which was important for the development of MITEs markers.

The 165.7Mb of passion fruit genome sequence was used to perform gene annotation with homologous species *Manihot esculenta* [17] and *Prunus persica* [18], and a total of 23,053 genes were predicted. The passion fruit genome size is 1,230 Mb [26], and the genome size involved in this study is approximately 13.5% of the total genome length. Therefore, we need to assemble the passion fruit genome to a higher level using high-throughput sequencing, especially at the chromosomal level, is particularly important.

By comparing the predicted protein sequences of passion fruit genome with the known protein sequences, we found that there were more genes related to carbohydrate metabolism, consistent with the fact that passion fruit is rich in sugar, fat, protein, vitamins and mineral elements [2,3]. However, there were less genes involved in environmental adaption in passion fruit genome, indicating that passion fruit may have poor capability to resist biological or non-biological stresses. At present, the main diseases of passion fruit are viral diseases, bacterial diseases and fungal diseases, among which fungal stem rot is particularly serious.

The comparison between passion fruit genome and the genomes of other eight species showed that only a few genes were unique in passion fruit. The unique family mainly contain genes of unkwnown functional proteins, retrovirus-related Pol polyprotein, zinc finger domain (CH2H2) proteins, and putative ribonuclease H protein. A number of genes are associated with retrovirus, which may suggest an important cause of the serious occurrence of viral disease in passion fruit. Specific regulatory sequences on DNA can bind to the corresponding regulatory proteins (transcription factors) and promote the initiation of transcription. In the unique family of passion fruit, the transcription factor family contains many genes, which may indicate that rich gene expression patterns are necessary for the continuous adaptation of passion fruit to the environment and to adjust its growth and metabolism.Moreover, in the evolutionary relationship, passion fruit is closer to *Populus trichocarpa* [23] and *Ricinus communis* [24], but the evolution rate is slower.

## ACKNOWLEDGMENTS

We thank Beltsville Agricultural Research Center. This work was supported by the Guangxi Natural Science Foundation of China (2018GXNSFBA281024, 2018GXNSFAA138124) and Guangxi Academy of Agricultural Sciences (2018YT19).

## DATA ACCESSIBILITY

All sequence data are gained from Genome Information for *Passiflora edulis* (BioProjects: PRJNA371406).

## AUTHOR CONTRIBUTIONS

X.H., Yang, Y.Y., Wu contributed to study design, Q.L., Tian, J.Y., Liu, Y.C., Huang, W.H., Huang contributed to data analysis, X.Z., Xia, H.F., Mou contributed to make tables and figures. All authors read and approve the paper.

## COMPETING INTERESTS

The authors declare that they have no competing interests.

